# Computational model of the human cochlea with motion of the layered osseous spiral lamina

**DOI:** 10.1101/2024.08.16.608342

**Authors:** Andrew A. Tubelli, Paul A. Secchia, Hideko Heidi Nakajima, Sunil Puria

## Abstract

**Purpose:** In the base of the human cochlea, the partition anatomy is distinct from the commonly recognized anatomy of laboratory animals. The human features a radially wide, osseous spiral lamina (OSL) and a soft-tissue bridge region that connects the OSL to the basilar membrane proper. In addition to the basilar membrane, the human OSL and bridge move considerably. We investigated the complex cochlear partition in human emphasizing the layered structure of the OSL with a finite element model. Model results were calibrated with experimental measurements of motion from optical coherence tomography.

**Methods:** The box model contained two fluid chambers separated by a cochlear partition and a helicotrema. Model geometrical and material properties either came from literature, measurements, or were tuned to produce a frequency-place map for the passive human cochlea as well as motion results similar to experimental measurements.

**Results:** The model motion results of the cochlear partition were similar to experimental results mostly within 5 dB but with differences at the high frequencies in both magnitude and phase beyond the best frequency. Around the best frequency location, the radial profile of cochlear partition motion was generally similar in both shape and magnitude. Sensitivity analysis, changing material-property parameters of the middle layer where the cochlear nerve fibers run between the layers of OSL plates, produced small changes in the model response and also showed negligible stress compared to the outer OSL plates.

**Conclusion:** These results suggest that the layered OSL anatomy is favorable as a conduit and protection for the nerve fibers while simultaneously functioning as a mechanical lever.

## INTRODUCTION

The anatomy and physiology of the basal cochlear partition in human is complex and different from laboratory animals [1]. In Figure 1, human cross-sectional cochlear histology (A) is compared to a stylized graphical representation of the partition (B). From the modiolus, two thin bony plates of the osseous spiral lamina (OSL) fan out laterally to the medial edge of the soft-tissue bridge. Auditory nerve fibers run between the bony plates of the OSL. From the spiral ligament of the lateral edge, collagen fibers project medially through the basilar membrane (BM), through the bridge, and insert onto the lateral edge of the OSL [2]. Altogether, this OSL-bridge-BM structure forms a partition that separates the cochlear fluid volume into the ducts of scala vestibuli and media on one side of the partition and the scala tympani on the other. The limbus has a roughly triangular profile and spans much of the OSL and part of the bridge. The organ of Corti abuts the BM and is separated from the OSL by the bridge.

**Fig. 1.**
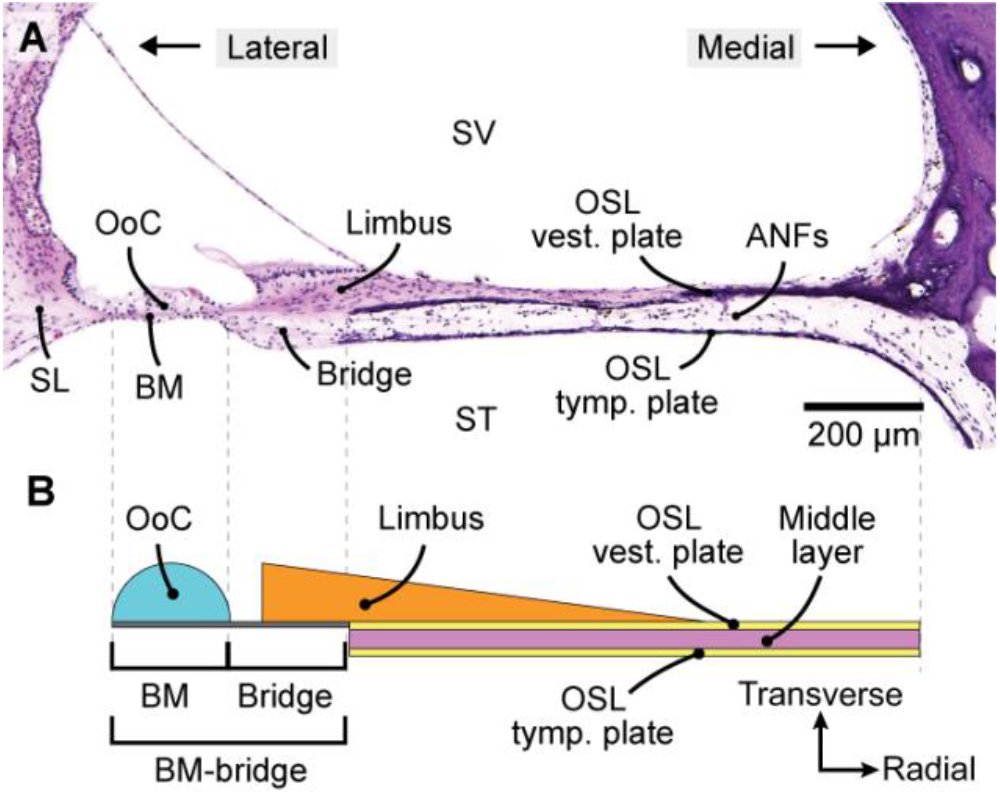
A: Labeled histological section of the human cochlear partition towards the base. B: Simplified graphical representation of the cochlear partition related to present model geometry. ANFs = auditory nerve fibers, BM = basilar membrane, OoC = organ of Corti, OSL = osseous spiral lamina, SL = spiral ligament, ST = scala tympani, SV = scala vestibuli, tymp. plate = tympanic plate, vest. plate = vestibular plate.

The OSL has a vestibular plate and a tympanic plate with a complex middle layer. Throughout the middle layer, thin bony pillars connect the vestibular and tympanic plates in a randomly distributed manner [2-5]. Some bony projections from the plates are not connecting pillars; rather, they resemble cave-like “stalactites” or “stalagmites”. Auditory nerve fibers course through the OSL middle layer radially between the bony cavernous structures from the modiolus towards the bridge and the organ of Corti. Thus, the OSL middle layer is a composite of bone and soft tissue.

Measurements of the human OSL show that the OSL is mobile in human [6]. Motion of the OSL in human cadaveric specimens was measured at the 2 kHz place [7] and 9-15 kHz place [1] with motion increasing from modiolus to the lateral aspect.

In addition to measurements, there has been work towards understanding the OSL through modeling efforts. Allaire et al. [8] predicted that the OSL would be mobile based on its cantilever structure and included it in an analytical model. Steele [9] considered the flexibility of the OSL to be important for high frequencies based on Rhode’s [10] measurements in the 7-kHz region in squirrel monkey. Raufer et al. [1] created a simple beam model that was consistent with experimental measurement of the radial profile of the cochlear partition.

Several finite element models of the human cochlea have been made. Most models are in the form of a box, i.e., a rectangular approximation of an uncoiled cochlea, or cochlear spiral and only account for the motion of the BM with rigid constraints on the medial and lateral ends, and sometimes soft tissues of the organ of Corti as well (e.g., [11-13]). Most human computational models have not included the OSL because they assumed that the OSL was rigid and thus has no motion. A few modeling studies have included an OSL. Koike et al. [14] added a single-layered OSL with width varying from base to apex and did not explicitly constrain its motion with fixation. However, in this model, the Young’s modulus of the OSL was made to be exceedingly stiff and thus effectively fixed with the intention to make it move very little. Borkowski et al. [15] included a single-layered mobile OSL with varying width and Young’s modulus from base to apex and favorably compares results to Stenfelt et al. [7]. Generally, computational models of the human cochlea have not included or considered the human structural anatomy, such as the complex multilayered structure of the OSL, the soft-tissue bridge, or the placement of the organ of Corti and limbus relative to these structures.

The aim of this study was to computationally model the human cochlea with more accurate and likely important anatomical complexities incorporated than in past models. A tapered box model of the human cochlea which emphasizes the mechanics of the layered OSL is presented. The human cochlear partition was modeled as a 3D fluid-filled cochlear duct, incorporating both anatomical findings and measurements of detailed cochlear partition motion. This model was tuned to be consistent with the cochlear frequency map and our OCT-derived measurements of the cochlear partition from the basal high-frequency region in fresh intact human cadaveric temporal bone specimens [16]. In situ cochlear partition images and motion were obtained through the intact round window membrane with OCT.

## METHODS

### Model development

#### Geometry

A tapered-box cochlear model with a partition separating the cochlear duct into two fluid chambers, one representing the scala vestibuli and scala media combined without Reissner’s membrane separating them, and the other representing scala tympani, was developed (Figure 2A). A two-chambered box structure is a common simplification for cochlear models that approximates the response of the spiral cochlea well while simplifying mathematical analysis [17]. The partition, shown in Figure 2B, included an OSL, a “BM-bridge” region (combination of BM and bridge labeled in Fig. 2B), limbus, and organ of Corti. Since the collagen fibers of the BM continue through the soft-tissue bridge region, both the bridge and BM are collectively labeled as “BM-bridge” in the present model. A helicotrema region was included at the apical end of the model by tapering off the partition to the modiolar side of the cochlea, leaving a triangular fluid connection between the two fluid chambers. Details about the geometrical dimensions of each model component follow below.

**Fig. 2.**
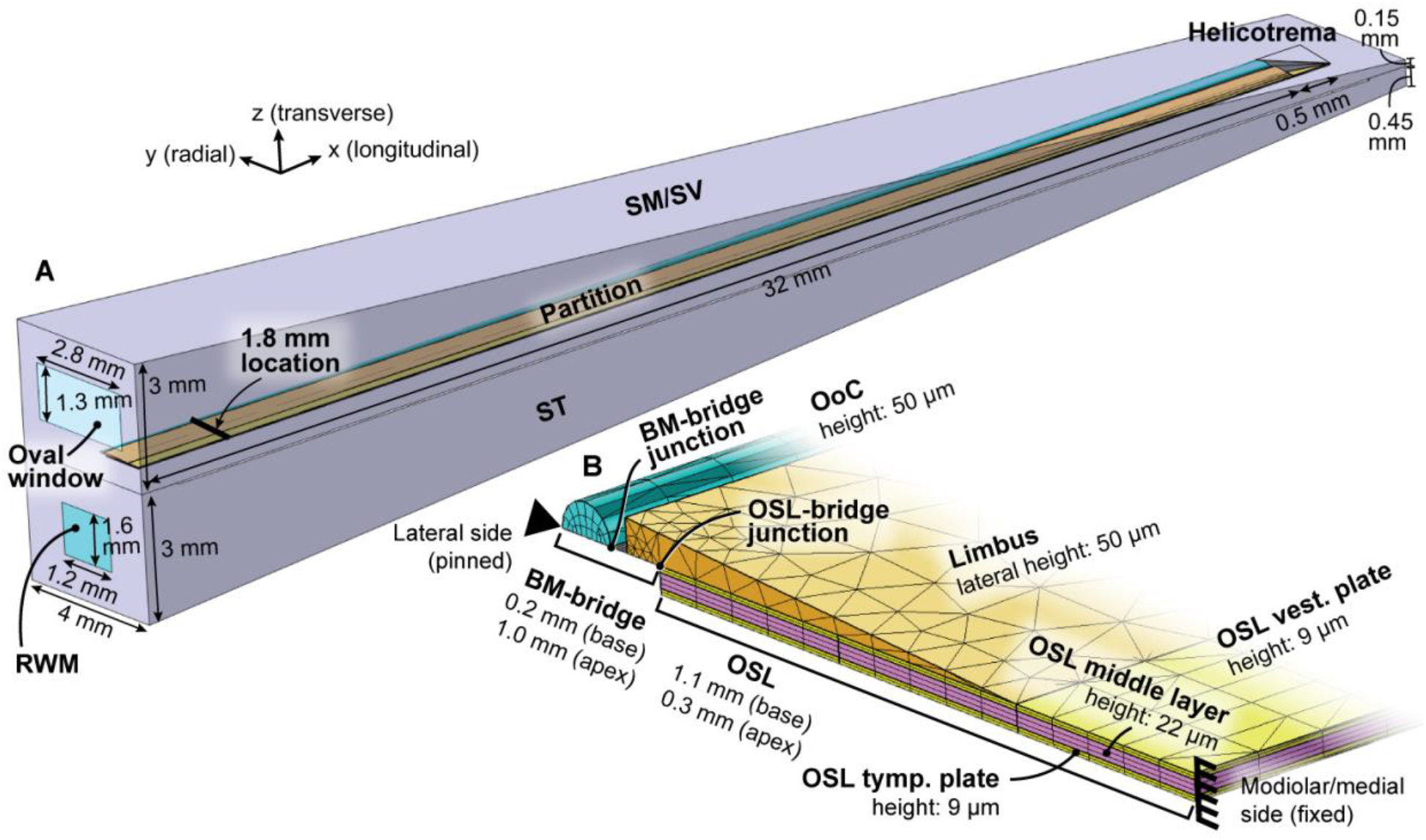
Model geometry and measurements. with a cross section of the cochlear partition at the base and boundary conditions. A: Full-length box model geometry. The model 1.8 mm measurement location is labeled. B: Scaled-up view of the basal-most partition with radial boundary conditions. OoC = organ of Corti, OSL = osseous spiral lamina, RWM = round window membrane, SM = scala media, ST = scala tympani, SV = scala vestibuli, tymp. plate = tympanic plate, vest. plate = vestibular plate.

Scalar cross-sectional areas at the base and apex were reported by Thorne et al. [18] and Wysocki [19]; however, since the model excludes the vestibule of the cochlea to instead form a geometrically simpler box, we needed to make accommodations for the windows. Ultimately the widths of both scalae were 4 mm throughout the length. The heights of both scalae at the base were 3 mm, and at the apex 0.15 mm for scala vestibuli and 0.45 mm for scala tympani.

The model partition was given a longitudinal length of 32 mm [20]. The radial widths of the OSL and BM-bridge as a function of longitudinal location were taken from histological measurements by Raufer et al. [2]. The BM and bridge had approximately equal widths along the length of the cochlea, both increasing from approximately 0.1 mm at the base to 0.5 mm at the apex. The OSL, on the other hand, has the opposite relation. The width decreased from 1.1 mm at the base to 0.3 mm at the apex. The total cochlear partition width (i.e., the sum of the OSL, bridge and BM) remained nearly constant at about 1.3 mm along the length of the cochlea.

The transverse height (i.e., thickness) of the OSL remained constant along the length of the cochlea. We looked at histological reconstructions to measure OSL plate thicknesses and total OSL thickness. On average, each plate had a thickness of approximately 9 μm but was variable within a range of 6-16 μm range both in radial and longitudinal directions. The model used individual plate thicknesses of 9 μm and a middle layer thickness (representing where the nerve fibers run) of 22 μm, totaling 40 μm in height (Figure 2).

The BM-bridge thickness varied from base to apex in the model which, along with the width, sets the stiffness gradient that produces a traveling wave and the cochlear place-frequency map. Even though Bhatt et al. [21] measured human BM thickness, such histological measurements could have inaccuracies. Alterations in dimensions of cochlear tissue occur by way of fixation, dehydration, or mechanical distortion from the cutting process, and such alterations differ across different types of tissue [22-24]. Thus, the BM-bridge thickness as a function of longitudinal position was the one geometrical parameter that was tuned to experimental data (see “Model tuning to experimental data” section). A piecewise linear function was ultimately used to describe the thickness (Figure 3). The BM-bridge thickness was set to 4 μm for the basal-most 8 mm, then decreased linearly to 0.45 μm at the helicotrema. Our thickness is only an estimate since the thickness of the bridge is different from the BM; however, the collagen fibers running through the BM-bridge likely dominate the mechanics. Average thickness data at four longitudinal locations from Bhatt et al. [21] is also plotted in Figure 3 showing that our estimate is within a factor of 2-3 of the experimentally measured thickness range.

**Fig. 3.**
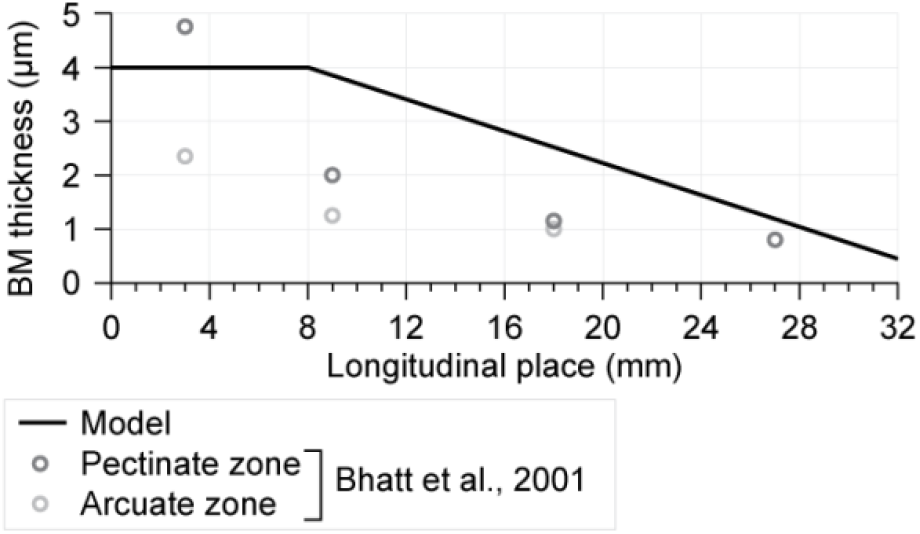
BM thickness in the model as a function of longitudinal place from base to apex.

Oval window area (3.64 mm^2^) and round window area (1.92 mm^2^) were calculated from an average of measurements from several literature sources [25-31]. Round window membrane thickness was uniform and approximated as 70 μm [32].

#### Material properties

The majority of material properties for the human cochlear structures have not been measured. Starting with parameters previously used by other modelers, the present model relied on sensitivity analysis in order to select key parameters that were then adjusted to produce results closest to the experimental data (see section “Model tuning to experimental data” for a more details on parameter sensitivity analysis). A summary of the material parameters is provided in Table 1. Tuned parameters are denoted with an asterisk.

**Table 1.**
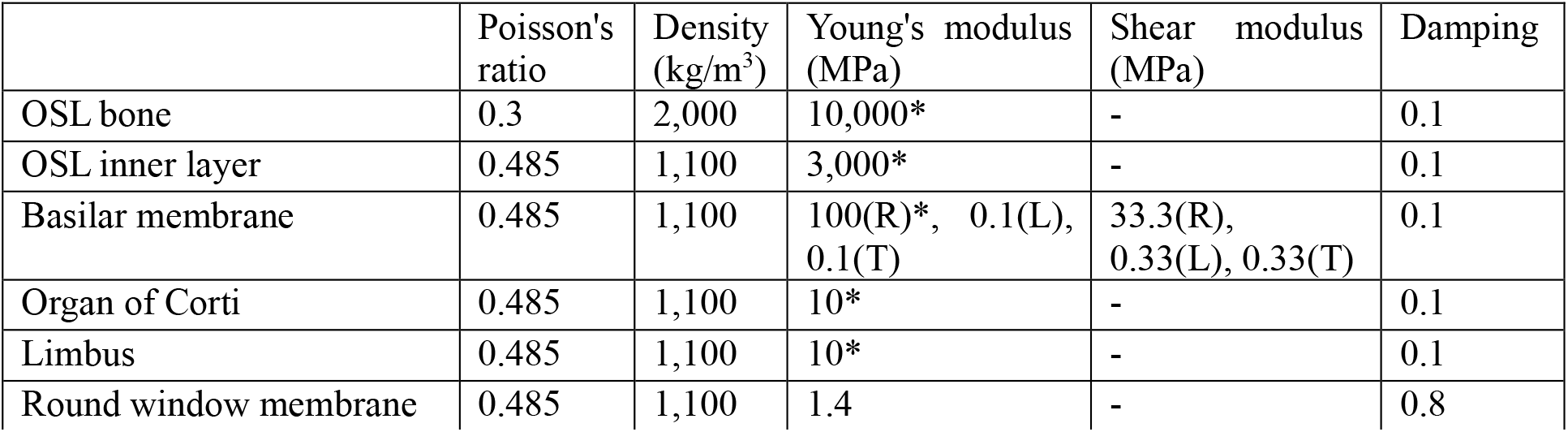
A: Labeled histological section of the human cochlear partition towards the base. B: Simplified graphical representation of the cochlear partition.

|  | Poisson's ratio | Density (kg/m<sup>3</sup>) | Young's modulus (MPa) | Shear modulus (MPa) | Damping |
| --- | --- | --- | --- | --- | --- |
| OSL bone | 0.3 | 2,000 | 10,000* | - | 0.1 |
| OSL inner layer | 0.485 | 1,100 | 3,000* | - | 0.1 |
| Basilar membrane | 0.485 | 1,100 | 100(R)*, 0.1(L), 0.1(T) | 33.3(R), 0.33(L), 0.33(T) | 0.1 |
| Organ of Corti | 0.485 | 1,100 | 10* | - | 0.1 |
| Limbus | 0.485 | 1,100 | 10* | - | 0.1 |
| Round window membrane | 0.485 | 1,100 | 1.4 | - | 0.8 |

Young’s modulus values varied among the modeled tissues: 10 GPa for OSL plates, 3 GPa for the OSL middle layer, 10 MPa for organ of Corti and limbus, and 1.4 MPa for the round window membrane.

The middle layer of the OSL representing nerve fibers and bony pillars was treated as having a composite Young’s modulus. Although not explicitly stated in Bom Braga [5], we estimated that the bony pillars could constitute 30% of the volume of the middle layer portion of the OSL. The OSL middle layer was therefore given a composite Young’s modulus value using the rule of mixtures for composite materials:

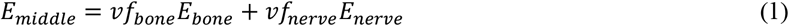

where E_bone_ is the Young’s modulus of bone given above (10 GPa), vf_bone_ is volume fraction of bone at 0.3, E_nerve_ is the Young’s modulus of nerve tissue taken as 10 MPa, vf_nerve_ is volume fraction of nerve tissue at 0.7. The composite middle layer Young’s modulus is ∼3 GPa. Note that because the bone Young’s modulus is much greater than that of the nerve tissue, Eq. (1) is approximately reduced to vf_bone_*E_bone_.

To investigate the contribution of stiffness of the middle layer, we performed sensitivity analysis by modifying the baseline model middle layer material. For the purposes of this sensitivity study, the baseline model is referred to as the “composite middle layer” model. Other variations of the model were investigated: in one model, the middle layer was given the same material as the bony plates (i.e., Young’s modulus, Poisson’s ratio, and density were made that of bone), effectively making the three OSL layers into a single thick bony plate. This model instance will be referred to here as the “bony middle layer” model. In another model, the Young’s modulus of the middle layer was reduced to 270 MPa, or 2.7% of the baseline value, which is in the range of biological soft tissues [33]. This instance will be referred to as the “soft-tissue middle layer” model.

The BM-bridge was treated as an orthotropic material, where the radial direction was stiffest since collagen fibers within the BM-bridge are primarily oriented radially. The Young’s modulus value of the radial direction was 100 MPa. This value did not change from base to apex; therefore, the model relied on the BM thickness as a function of length as the parameter to set the stiffness gradient in the model that produced the traveling wave, The longitudinal and transverse direction values were lower than the radial direction by three orders of magnitude (0.1 MPa), consistent with a very soft ground substance that reduces longitudinal coupling [34].

While we don’t know the Young’s modulus of the BM-bridge for certain, we know that the BM-bridge is one of the stiffest soft tissues in the partition due to the presence of collagen fibers [35-37], which are typically in the range of 1-20 GPa [38,39]. The overall stiffness of the BM-bridge depends on the volume fraction of collagen fibers. Although we did not specify volume fraction in this model, we can calculate volume fraction from our chosen material property values using the rule of mixtures for composite materials:

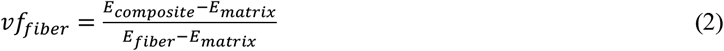

Here vf_fiber_ is the fiber volume fraction (ratio) of collagen, and E is Young’s modulus. Given E_composite_ of 100 MPa, E_matrix_ of 0.1 MPa (that of the BM-bridge transverse and longitudinal directions) and E_fiber_ within the range of values for collagen, the volume fraction is estimated to be 0.5-10%. This is close to the range of volume fractions from base to apex calculated by Kapuria et al. [40] for guinea pig (1-8%) and gerbil (0.8-16%).

Since the BM-bridge was orthotropic, the shear modulus was also specified in the model. The relation between shear modulus and the other mechanical properties for an orthotropic material is complex and not well understood; therefore, for simplicity, each individual orthogonal direction was given a shear modulus value equal to one third of the respective Young’s modulus value, as has been done in other modeling studies (e.g., [41]).

Structural damping of all model components, with the exception of the round window membrane, was set to 0.1 (e.g., [41]). Damping of the round window membrane was set to 0.8. The higher damping at the round window is designed to compensate for radiation impedance of the middle ear cavity.

Density of bone was set to 2,000 kg/m^3^, and density of all soft tissue components was set to 1,100 kg/m^3^, similar to other auditory models [42,43].

Poisson’s ratio of bone (the OSL plates) was set to 0.3, a typical value used in modeling bone (e.g., [44,45]), and all soft tissue components were assumed close to incompressible at 0.485 (e.g., [41]).

Cochlear fluid was modeled as water for simplicity.

#### Physics and boundary conditions

Three aspects of physics were combined in this model: thermoacoustics for the fluid domains (i.e., scala vestibuli and scala tympani), solid mechanics for the three layers of the OSL, limbus and organ of Corti, and shell mechanics for the BM-bridge and round window membrane. Thermoacoustics is used for its inclusion of fluid viscosity. Shell mechanics is a computationally efficient approximation to model structures like the BM-bridge and round window membrane that are transversely thin in comparison to other dimensions.

The partition was given a fixed boundary condition on the medial side of the OSL. On the lateral end of the BM-bridge (lateral end of BM), the applied boundary was given a pinned condition corresponding to fixed translation and free rotation (Figure 2B).

The normal vectors of the BM-bridge shell project towards the scala tympani so that the top of the BM-bridge is level with the top of the OSL vestibular plate.

Input velocity stimulus at the oval window was constant across frequencies at 4 μm/s. All model results, as with experimental results, are reported as displacements relative to stapes input. For the model, input velocity was converted to displacement for these calculations. The model is linear, and experiments have also confirmed linearity.

#### Finite element model details

The model was created using COMSOL Multiphysics (www.comsol.com) version 6.1. The final mesh was composed of the 469,919 elements consisting of 296,946 tetrahedra, 31,925 pyramids, 57,158 prisms, and 83,890 hexahedra. There were 3,962,263 degrees of freedom in total. The computer used was a Dell desktop running a Linux operating system with two 20-core Intel Xeon Processors, 40 cores in total, and 512 GB of RAM. With this hardware, the simulation time was approximately 30 minutes per frequency. The frequencies used in the model were between 3-14 kHz at one-third octave intervals with additional frequencies added after 10 kHz. In total, the model was solved at 11 frequencies.

### OCT measurement methods

Cochlear partition transverse motion in response to sound presented to the ear canal was measured while visualizing structures with OCT vibrometry through the round window membrane on a fresh cadaveric human specimen (16 hours postmortem). Multiple radial location measurements near the scala tympani surface of the cochlear partition were made. Cross-sectional imaging and vibrometry measurements of the cochlear partition were made using a Spectral-Domain OCT system with a 900-nm center wavelength and A line-scan camera frame rate of 46-kHz (GAN620C1, Thorlabs, Germany). The imaging axial resolution was 2.23 µm (in water) and the lateral resolution approximately 8 µm, using a 36 mm, 0.055NA, 2x objective lens. Custom-built LabVIEW-based software (VibOCT versions 2.1.5) recorded images and made vibration measurements. SyncAv (version 0.47) generated pure-tone sequences from 0.1-21 kHz ranging from ∼80 to 110 dB in the ear canal where linearity was confirmed. The measurement noise floor was 0.3 nm or better above 2 kHz and SNR criteria of 6 dB was used for all measurements reported. Stapes velocity was also measured with laser Doppler vibrometry (CLV-3D, Polytec, Germany). OCT displacement measurements within the cochlear partition were referenced to the stapes displacement.

### Model tuning to experimental data

The parameters chosen for model tuning were those that were not well known in the literature or those that produced significant changes in the model response during single-parameter sensitivity studies. The tuned parameters were BM-bridge thickness as a function of length, and Young’s moduli of the three OSL layers, BM-bridge, organ of Corti, and limbus.

To quantitatively assess how well the model performed with parameter variations, we calculated the difference between the model and respective experimental measurements: the frequency-place map, displacement frequency response normalized to oval window input, and radial profile of displacement relative to input. By randomizing unknown material properties within physiological limits, we converged on model parameters that produced the closest match to the measurements, i.e., the minimization of differences between model and data for all measurement types simultaneously. Minimization criteria were partly quantitative in determining the difference between model and data response values at each frequency, and partly qualitative in looking at the difference in response shapes. The best set of parameters used in the model satisfied the minimization criteria but may not be unique.

The Greenwood cochlear frequency map [46] for human was used. Since the model and all human experimental data were passive, the Greenwood map was shifted one half octave to the base to account for reduction of the CF in the passive cochlea [47].

Both the frequency response and radial (spatial) profile of motion responses were compared between the model data and experimental vibrometry measurements. Two key anatomical locations, the OSL-bridge junction and the BM-bridge junction (these junctions are labeled in Figure 2B), were used to compare frequency responses between experimental and model results. These junction points will be referred to as the OSL and BM, respectively, in the discussion of results going forward. Points along the scala tympani surface of the cochlear partition were used to compare radial motion responses.

## RESULTS

### Baseline model tuning

#### Cochlear map

Most individual frequencies in the frequency-place map of the model produced a peak response longitudinally within ±1 mm of their expected locations from the Greenwood map that was shifted to represent the passive cochlea (Figure 4).

**Fig. 4.**
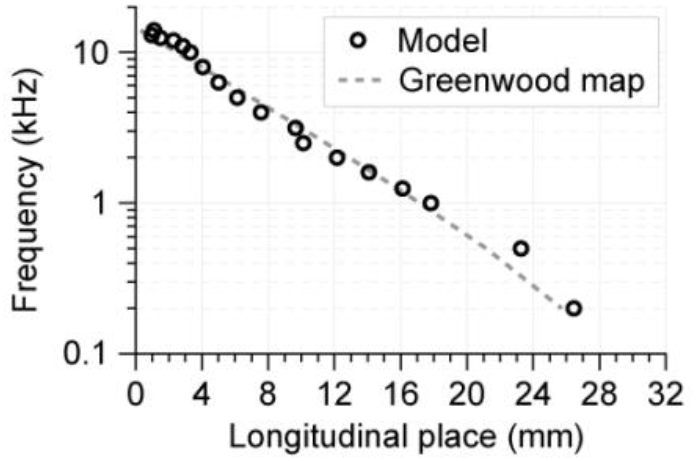
Tuned-model frequency-place map vs. Greenwood map [46]. The Greenwood map is shifted basally a half octave to mimic a passive cochlea.

#### OSL and BM frequency response

The measured and model displacement frequency responses of the BM-bridge junction and the OSL-bridge junction near the scala tympani surface at the cochlear base are shown in Figure 5. The model results were taken at 1.8 mm from the base. For the experimental measurement, longitudinal location was estimated to be approximately 1 mm from the base. The experimental location area estimation was broadly based on the best frequency (BF) of 12.5 kHz, the passive Greenwood map, and a comparable cochlear length to the model. Then, because we estimated the cochlear length in the experimental specimen, the location for the model was finely chosen so that results were closest to experimental data, resulting in a slightly different BF of 11 kHz for the model.

**Fig. 5.**
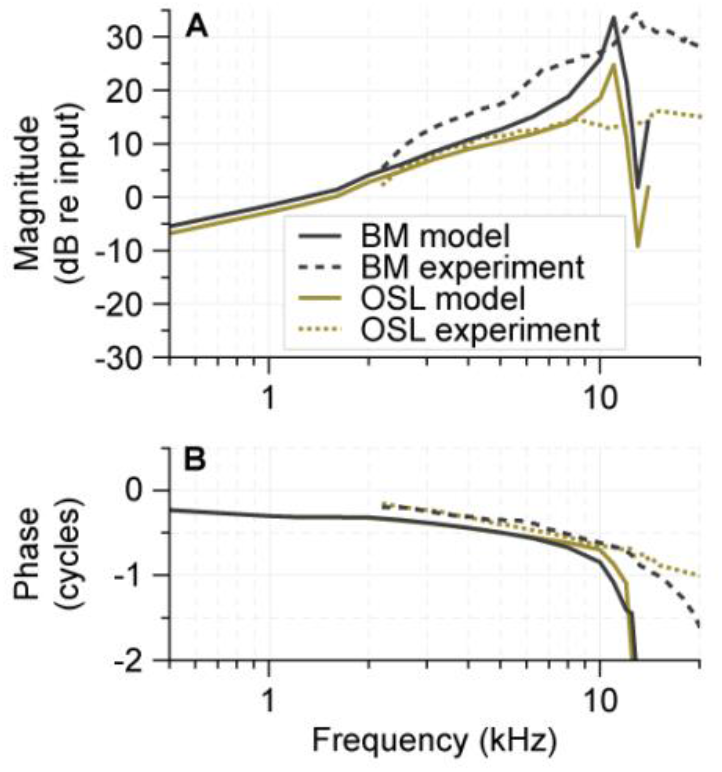
Comparison of motion frequency response at the base for the tuned model and the experimental measurements at the BM-bridge junction (BM) and OSL-bridge junction (OSL). Experimental data shown are output displacements normalized to stapes displacement, interpolated at the model frequencies. The longitudinal place was about 1 mm for the experimental data and 1.8 mm for the model. The model result shown is displacement normalized to oval window displacement. The BM lines correspond to the point of maximum motion, which in the model is the BM-bridge junction, and in the experiment was approximately 20 μm lateral to the BM-bridge junction.

The experimental data had similar trends to the model results with some differences. For most frequencies, the magnitude of the model response was within 5 dB of the experimental response up to the BF peak for the model (Figure 5A). In the model, the magnitudes of OSL and BM rose proportional to frequency (+6 dB/oct) from 0.5 kHz to about ½BF (6 kHz), with a slightly shallower slope below about 2 kHz. With increasing frequency, both model OSL and BM magnitudes reached their peak at BF with the BM being 9 dB higher than the OSL. At higher frequencies, both model OSL and BM magnitudes dropped sharply after the peak frequency. As a comparison, the experimental measurement (dashed lines in Figure 5) magnitudes at BF were slightly higher and more broadly peaked in the BM than in the model. The difference in magnitude between OSL and BM in the experimental measurements was also larger than in the model results.

The phases for the model and experimental data are plotted in Figure 5B. For the model, both BM and OSL phases were similar and decreased from approximately −0.25 cycles at 0.5 kHz to −0.5 cycles at ½BF, above which the phases decreased rapidly. This type of phase response is consistent with a fast traveling wave (low group delay) in the low frequency region and slow traveling wave (fast group delay) as the input frequency increases above about ½BF [48]. For the experimentally measured OSL and BM, the phase frequency responses were nearly identical up to BF; both OSL and BM phases decreased with frequency and were generally similar to the model phases to about ½BF. However, the model phases were generally more negative than the experimental phases by about 0.1 cycles for frequencies less than ½BF. One important deviation is that at frequencies ½BF and higher, the modeled phases decreased rapidly with frequency while the experimental phases did not. Reasons for this are not clear but one possible reason might be that the stiffness gradient in the model might be steeper in comparison to that in the measured specimen.

#### OSL and BM radial profile

Another perspective of the cochlear base is to look at the cochlear partition’s radial profile, focusing on the transverse displacement at varying radial locations. Figure 6 shows the model and experimental cochlear partition transverse displacement normalized to stapes input displacement, plotted against radial location. The three plots are for three different frequencies: (A, B) best frequency (11 kHz in the model, 12.5 kHz in the experiment), (C, D) 8 kHz, and (E, F) 4 kHz.

**Fig. 6.**
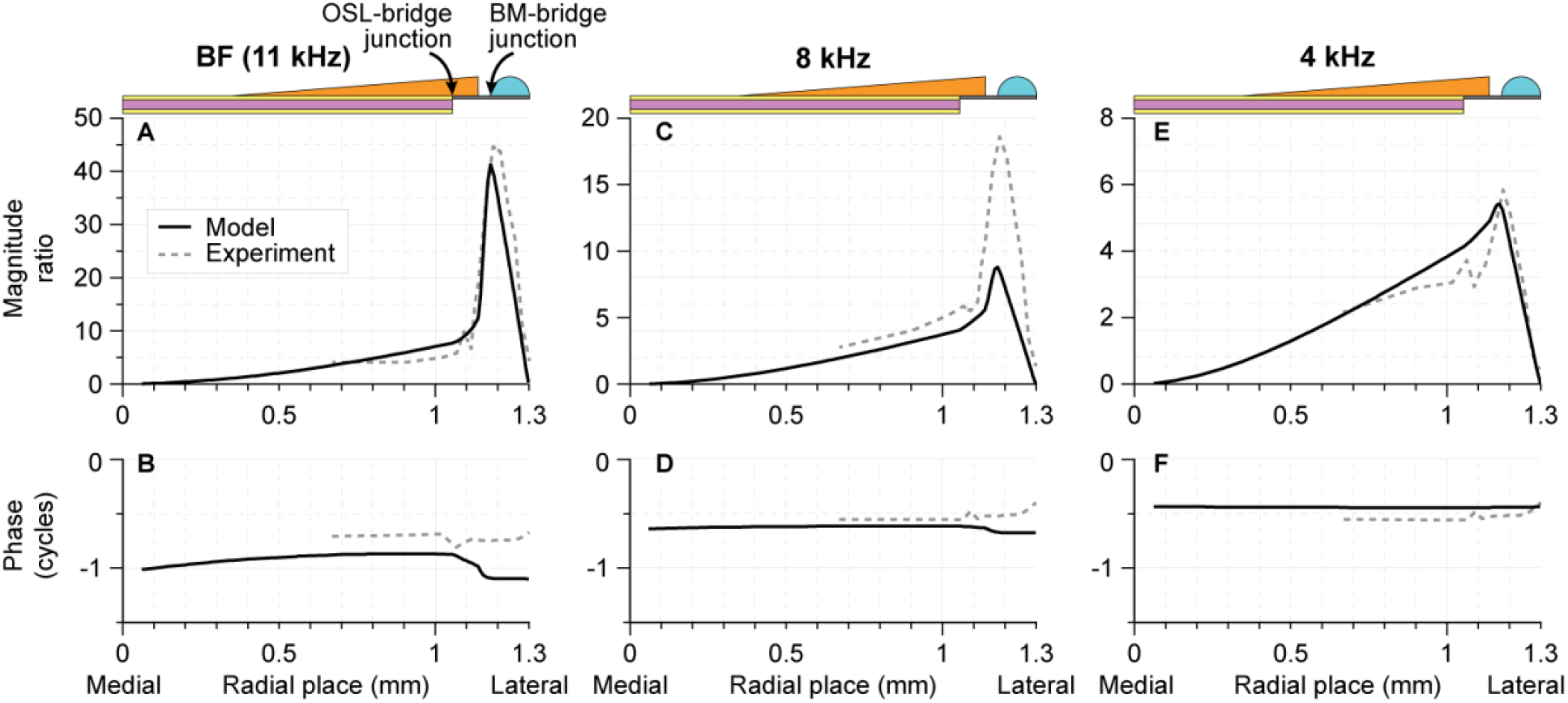
Comparison of the model and OCT data for transverse motion referenced to input motion of the stapes, plotted against radial place. Experimental data shown are transverse displacements normalized to stapes displacement. The model results shown are transverse displacements normalized to input displacement. A, B: both experimental and model data lines correspond to their respective best frequencies (11 kHz in the model, 12.5 kHz in the experiment); C, D: model frequency of 8 kHz, experimental frequency of 8.114 kHz; E, F: model frequency of 4 kHz, experimental frequency of 3.93 kHz. The longitudinal location was at the base, where the model was at 1.8 mm and the experiment was at 1 mm from the basal end. Note the differing y-axis scales.

Both model and experimental OSL increased in magnitude with radial position from the medial end to the OSL-bridge junction. Then further laterally, past the limbus and closer to the border of the BM-bridge junction, the motion increased more rapidly, reached a peak and then decreased. In the model, the peak occurred directly at the BM-bridge junction (also where the foot of the inner pillar cells would have been), while for the experimental measurements the peak occurred more laterally at the BM. This subtle difference in peak radial locations can possibly be attributed to the absence of relatively stiff pillar cells and the tunnel of Corti in the model, which were likely present in the measured temporal bone. The tunnel of Corti can stiffen the BM causing the peak to occur more around the outer pillar [9].

At BF (Figure 6A) and 4 kHz (Figure 6E), the radial profile of the magnitudes between model and experiment were mostly similar. For 8 kHz (Figure 6C), the OSL magnitude of the model was similar to the experiment, but it is unclear why the BM magnitude peak of the model was only half that of the experiment.

The phases at each frequency (Figure 6B, D, F) of the model were close to experimental phases, within a quarter cycle. At higher frequencies (8 kHz and 11 kHz), a decrease in phase can be seen in the bridge region between the OSL-bridge and BM-bridge junctions. This feature is also present in some of the LDV experimental results in Raufer et al. [1].

### The layered OSL

A major thrust of this study was to explore the role of the mobile OSL in the overall cochlear partition mechanics, and in particular the role of the middle layer where the auditory neurons traverse between the thin bony vestibular and tympanic plates. Towards this end, aside from the baseline composite middle layer model, we also simulated a “bony middle layer” model of 100% bone and a “soft-tissue middle layer” model with a reduced Young’s modulus.

Figure 7 shows the frequency responses of the BM and OSL for all three middle layer Young’s modulus variations, both magnitude (A, C) and phase (B, D). The radial profiles of motion for the three model variations are in Figure 8. As shown in both Figure 7 and Figure 8, comparisons across the three models resulted in similar BM and OSL motion across frequencies and across radial locations, though small (within 10 dB and fractions of a cycle) differences from the composite baseline model were seen. Frequency-place maps were similar for the three models (not shown).

**Fig. 7.**
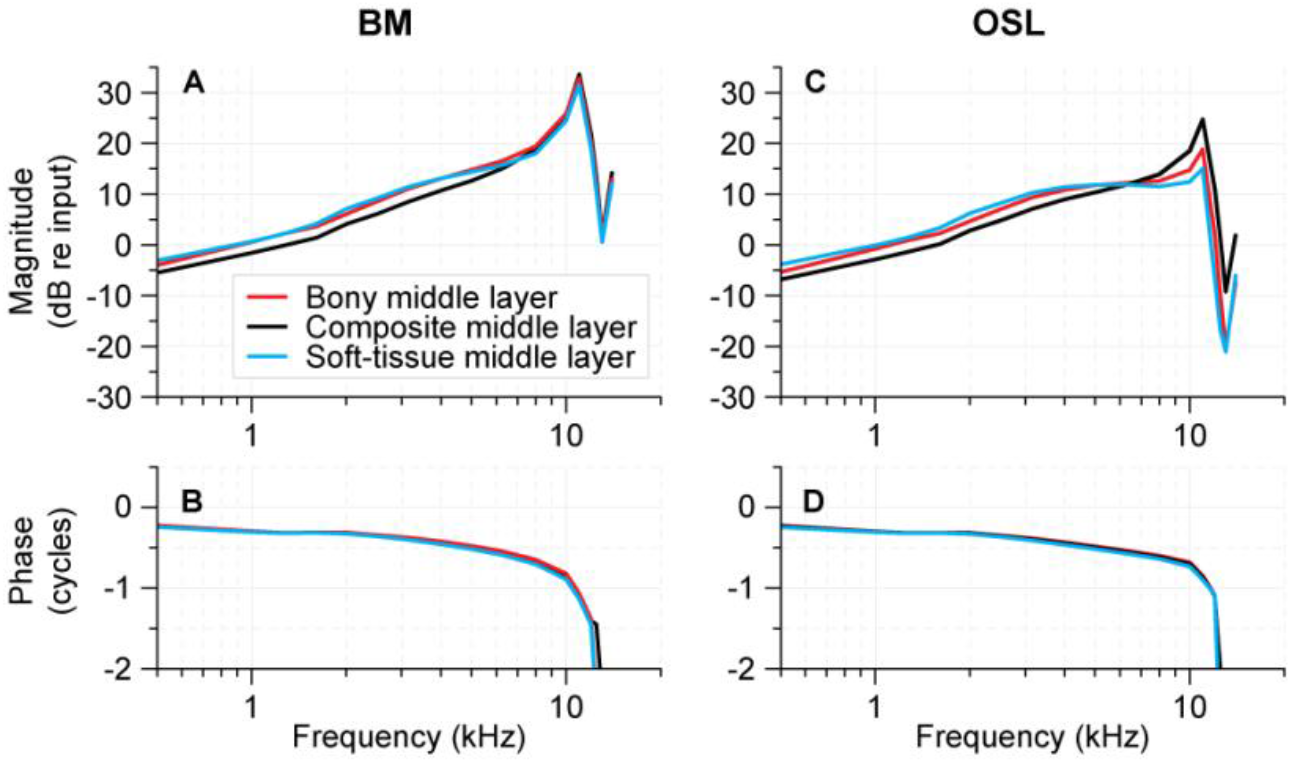
The frequency responses for the transverse displacement of the BM (left) and OSL (right) referenced to input displacement. The OSL middle layer Young’s modulus was varied for the three different model variations: bony middle layer model (10 GPa) similar to bone of upper and lower plates (red); the composite middle layer model (3 GPa, baseline) representing a composite of bone and soft tissue (black); and the soft-tissue middle layer model (270 MPa) representing a soft tissue at 3% of the baseline composite value (blue).

**Fig. 8.**
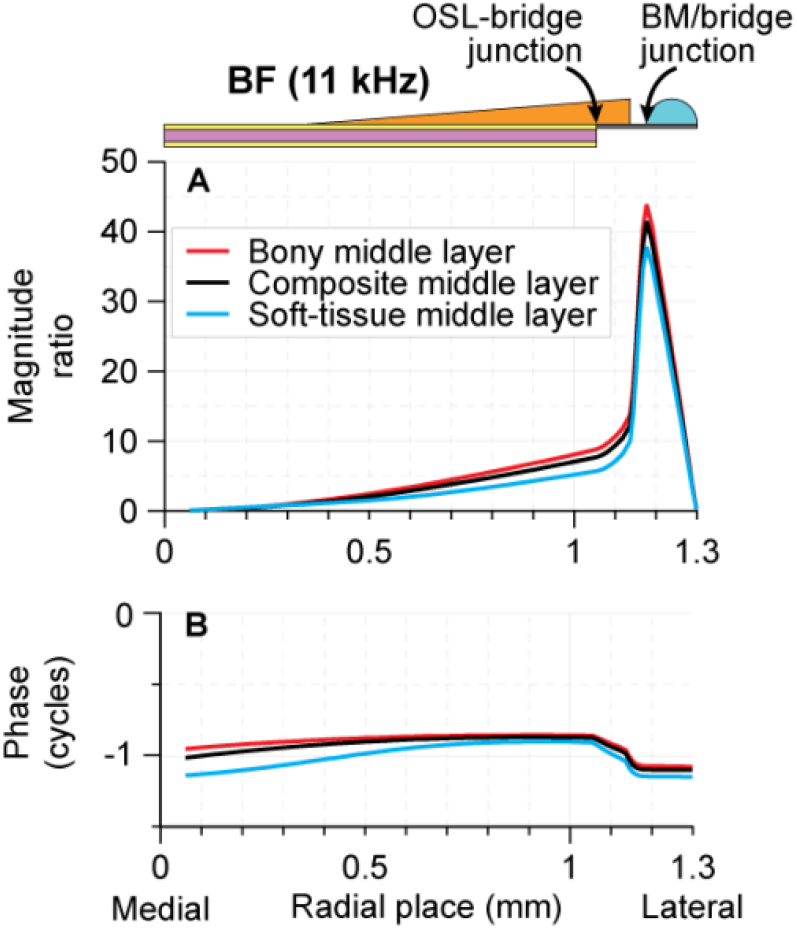
The transverse motion referenced to input displacement at different radial points. The OSL middle layer Young’s modulus was varied as described in Figure 7 caption. The longitudinal place was 1.8 mm with a BF of 11 kHz.

#### Stress in the OSL layers

The result that the Young’s modulus of the middle layer could be changed by a factor about 30 without significantly affecting the motion of the OSL and BM was surprising. We thus investigated the effect of the middle layer further by looking at the stress within it. In mechanics, one can look at several different types of stresses.

The von Mises stress (σ_v_) is a measure that gives an overview of multiaxial stress in a structure as a scalar value. The von Mises stress in the baseline model is plotted at 1.8 mm from the base in Figure 9. The phase of the motion cycle plotted for each frequency was chosen to correspond to the maximum stress, and color encodes the stress at that phase. It turned out that the maximum von Mises stress was at the maximum deflection of the OSL-bridge junction. The model also shows that stress is confined to the OSL plates while the stress in the OSL middle layer, limbus, and OoC were relatively insignificant. The composite, bony, and soft-tissue middle layer models all had similar profiles where maximum stress occurred in the OSL plates (model results not shown for bony and soft-tissue model variations).

**Fig. 9.**
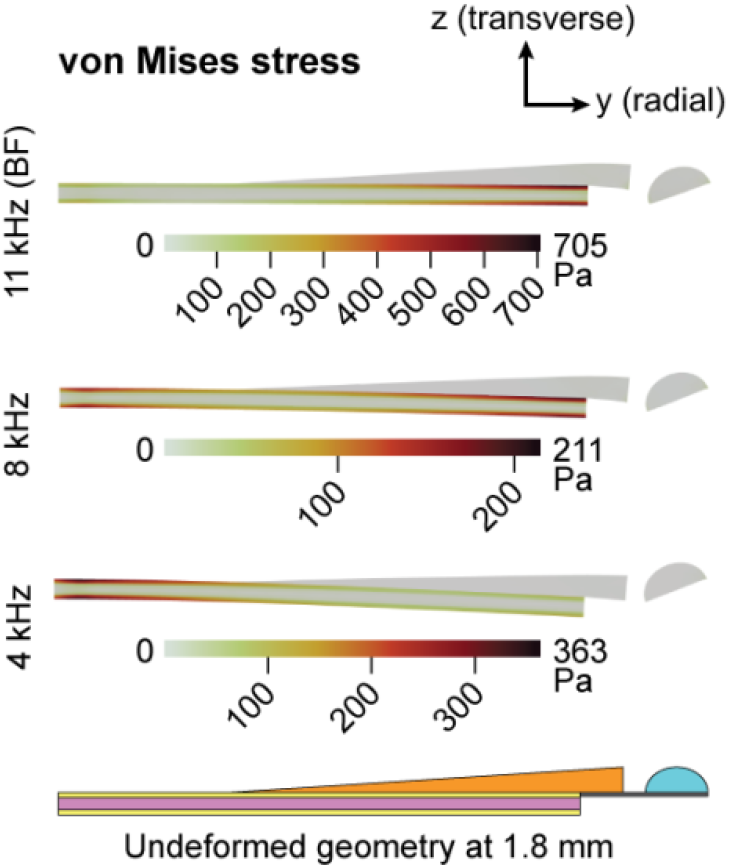
The von Mises stress in Pa for the baseline model at 1.8 mm from the base for 11 kHz (BF), 8 kHz, and 4 kHz. Note, each frequency has a separate corresponding scale. Color denotes the maximum stress experienced at each point in the structure at the phase of displacement corresponding to the maximum downward deflection of the OSL tip. Deformation is scaled for visibility (36,000x at 11 kHz, 75,000x at 8 kHz, 171,000x at 4 kHz). The undeformed graphical representation of the partition cross section at 1.8 mm is shown at the bottom of the figure. The majority of the nonzero stress is situated in the OSL plates.

While the von Mises stress gives an overall measure of the different stresses, another way to look at stress in a system is through the second Piola-Kirchhoff (2^nd^ PK) stress tensor. The 2^nd^ PK stress has normal (σ_x,y,z_) and shear (τ_xy,xy,yz_) stress components. Figure 10 shows the three highest stresses of the baseline model, namely normal radial stress σ_y_, normal longitudinal stress σ_x_, and shear stress τ_xy_ in the radial-longitudinal plane of the OSL. The phase of the deflection cycle plotted for each frequency was chosen such that it corresponds to the maximum positive stress. Note that unlike the von Mises stress, the phase corresponding to maximum 2^nd^ PK stress components in most cases do not correspond to the phase at maximum deflection. The model shows that all stress components, regardless of frequency, were localized to the OSL plates while the stresses in the OSL middle layer, limbus, and OoC were relatively insignificant. The opposite polarity of stress can also be appreciated with the 2^nd^ PK components, denoted by color, where positive normal stress denotes tension. The middle layer of the OSL remains stress-neutral in the model at all frequencies.

**Fig. 10.**
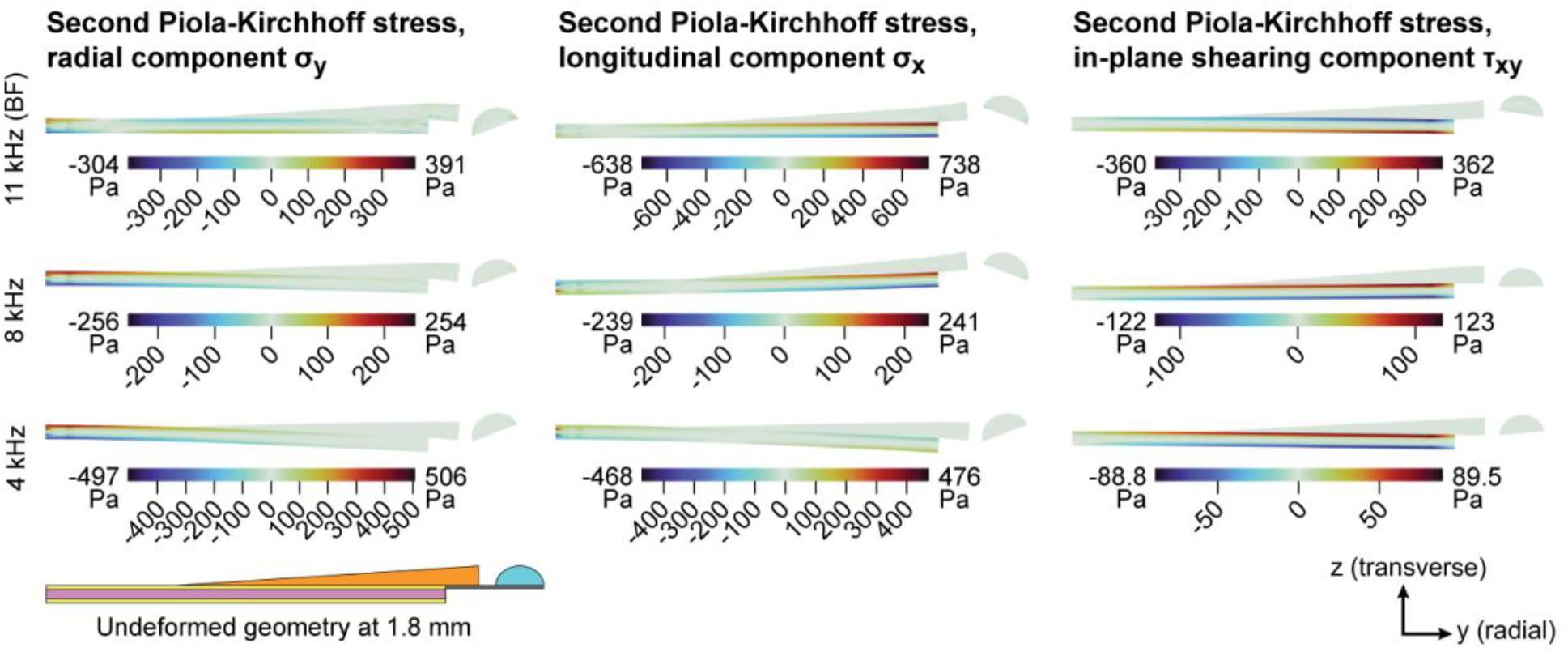
The Second Piola-Kirchhoff stress in Pa for the baseline model at 1.8 mm from the base for 11 kHz (BF), 8 kHz, and 4 kHz. Each frequency and component have a separate corresponding scale. Color denotes the maximum stress experienced at each point in the structure at a phase of displacement corresponding to greatest magnitude of positive stress in the deformation cycle. Deformation is scaled for visibility (36,000x at 11 kHz, 75,000x at 8 kHz, 171,000x at 4 kHz). The majority of the nonzero stress is situated in the OSL plates.

## DISCUSSION

The goal of this study was to develop a finite element model of the human cochlea which incorporated the anatomy of human, specifically the multilayered OSL and the presence of a soft-tissue bridge. The study further investigated the mechanics of the layered OSL encompassing two thin outer bony plates and its cavernous middle layer. The model was built using a combination of histology and OCT imaging measurements of the human cochlear partition. As is the case for many finite element modeling approaches, determining material properties of anatomical structures was a challenge that required parameter tuning. The chosen set of parameters were such that the model results were generally consistent with OCT-based anatomical and vibrometry measurements of the human cochlear partition (BM and OSL displacements as a function of frequency for a basal high-frequency location, as well as displacements at various radial locations at a few frequencies). Also, the model was consistent with the cochlear frequency-place map from the literature. The model results were generally consistent with experimental measurements, with some differences discussed below. Further analysis of stresses within the cochlear partition predicted that the middle layer of the OSL, where the neutral plane resides, is an optimal location of the auditory nerve for protection against stimulation or damage during OSL movement.

While there was good overall agreement between model calculations and measurements, there are noteworthy differences that merit discussion.

The frequency response in the model is more sharply peaked at BF than in the experimental measurements for both the BM and OSL (Figure 5A). The experimental signal was very low at high frequencies approximately above 10 kHz, resulting in lower signal-to-noise. In the experiment, the OSL motion at BF is 20 dB lower than the BM, suggesting worse signal-to-noise for the OSL and might explain why the experimental OSL data lacks a clear peak altogether.

In the experimental measurements, the magnitudes do not have sharp peaks and the phases do not drop off as quickly as the model results, suggesting the possibility that there is more damping in the experiments than incorporated into the model. This could be because the model did not incorporate the microstructures and fluid spaces of the organ of Corti. For example, the model did not contain a tectorial membrane, and thus did not include a sub-tectorial fluid gap. Incorporation of such structure and fluid spaces is expected to result in greater damping in the organ of Corti [49-51].

We present experimental data measured in one fresh healthy-looking human specimen. Because there is mechanical variability across human ears, our modeling here has limitations.

### Structure and function of the layered OSL

The middle layer of the OSL provides a passage for nerve fibers. The nerves travel medially from the hair cell synaptic terminals in the OoC, through the habenula perforata, through the bridge, then through the middle OSL layer, and into Rosenthal’s canal [20]. Within the middle layer, there are bony pillars that connect the two plates, but their columnar structure keeps the plates separated as visualized by Bom Braga et al. [5]. The cavernous non-bony space serves as a network of passages for the nerve fibers to go through. With a mobile OSL, deflection of the plates could possibly traumatize the nerves by introducing stress in the auditory nerve fibers. Auditory nerves can respond to mechanical stimulation and could produce firing that does not originate from the inner hair cells [52]. The three-layered OSL construction is possibly optimized for minimal mechanical stimulation of nerve fibers.

Changing the Young’s modulus of the middle layer of the OSL by nearly 30 times such that the entire OSL can be approximated as 100% solid bone at one extreme (the bony model variation of this study), or a layered OSL with a soft-tissue middle layer at the other extreme (the soft-tissue model variation), resulted in small differences in response of the OSL and BM motion (Figures 7 and 8). This means that, while the general layered structure of the OSL may be anatomically important, the material (bony or soft-tissue) in the middle layer of the OSL had minimal impact on overall OSL motion.

The mechanics of beam theory give insight to this result. The central portion of a symmetric cantilever beam forms a neutral plane with zero stress compared to the outer portions that bear the load. Our result of the overall motion being very similar whether the middle layer is made of bone or soft tissue is consistent with this theory. When stress is applied to a beam, there is compression on one side and expansion on the other with a neutral plane in between, parallel to the cross section of the beam and perpendicular to the plane of bending (illustrated in the stress results for the radial component in Figure 10). The middle layer of the OSL contains the neutral plane which runs from the base to the apex. The partition cross section is not entirely symmetric due to the limbus situated adjacent to the vestibular plate; nevertheless, the neutral plane is still visualized to be between the OSL plates.

Because there is no stress along the neutral plane, the material used to construct it will have little consequence in comparison to the material properties of the upper and lower plates. This is very similar to the design of an I-beam in construction materials where the middle portion of the beam is removed to save material cost or lower weight without sacrificing structural integrity. This type of design would optimally allow the auditory nerve fibers to pass through the OSL middle layer and into Rosenthal’s canal while minimizing stress to the nerve fibers. In short, what we see is localization of compressive and tensile stress to the OSL plates with little to no stress in the middle OSL layer where the auditory nerve fibers reside and thus potential avoidance of nerve stimulation.

In this study’s tapered box model with Cartesian coordinates, we have made the OSL without curvature for convenience and computational economy. In the very base of the cochlea, the OSL is seldom so straight as in Figure 1A. In our model, the OSL is symmetric and therefore the neutral plane is in the middle. But in the very base of human cochlea, at the hook, the OSL is a curved structure, thus, that neutral plane shifts towards the inner radius of the bend. We did not explore curvature of the OSL, but we suspect that the difference in Young’s modulus between the outer bony plates and the middle layer will still result in stresses being localized at the plates.

The motion of the OSL plates is primarily in the transverse direction with a longitudinally propagating travelling wave. This traveling wave was shown to result in high stress in the longitudinal direction of the bony OSL-bridge junction, especially at BF (Fig. 10, middle column showing σ_x_).

The OSL plates are described as being porous [2,5]. We did not explore porosity in our model because of the increase in computational cost and the assumption that it is not a major factor in understanding OSL mechanics. Given that our model was tuned, and given a porosity around 50% [5] we would predict that we would need a higher Young’s modulus of bone by a factor of two with porosity implemented to achieve the same model tuning.

## CONCLUSIONS

At the middle layer of the OSL, the cavernous structure with bony pillars may be important for creating a non-traumatizing passage for the auditory nerve fibers. The nerves would pass through the middle layer along a neutral plane free from mechanical stress while the stiff vestibular and tympanic plates of the OSL take on the stress-bearing load.

## ACKNOWLEDGMENTS

The authors thank Ahsan Cheema, John Guinan, Yanli Wang, John Zhang, and Aleks Zosuls for their helpful discussions. Funding for this study was provided by grant R01 DC013303 from the NIDCD of NIH.

## STATEMENTS AND DECLARATIONS

The authors have no competing interests.

## REFERENCES

[1] Raufer S, Guinan JJ, Nakajima HH (2019) Cochlear partition anatomy and motion in humans differ from the classic view of mammals. Proc Natl Acad Sci USA 116:13977–13982. 10.1073/pnas.1900787116

[2] Raufer S, Idoff C, Zosuls A, et al (2020) Anatomy of the Human Osseous Spiral Lamina and Cochlear Partition Bridge: Relevance for Cochlear Partition Motion. JARO 21:171–182. 10.1007/s10162-020-00748-1

[3] Fleischer G (1973) Studien am Skelett des Gehörorgans der Säugetiere, einschließlich des Menschen. Säugetierkundliche Mitteilungen 21:131–239.

[4] Rask-Andersen H, Liu W, Erixon E, et al (2012) Human Cochlea: Anatomical Characteristics and their Relevance for Cochlear Implantation. The Anatomical Record 295:1791–1811. 10.1002/ar.22599

[5] Bom Braga GO, Parrilli A, Zboray R, et al (2023) Quantitative Evaluation of the 3D Anatomy of the Human Osseous Spiral Lamina Using MicroCT. JARO 24:441–452. 10.1007/s10162-023-00904-3

[6] Kohllöffel LUE (1983) Problems in Aural Sound Conduction. In: de Boer E, Viergever MA (eds) Mechanics of Hearing. Springer Netherlands, Dordrecht, pp 211–217.

[7] Stenfelt S, Puria S, Hato N, Goode RL (2003) Basilar membrane and osseous spiral lamina motion in human cadavers with air and bone conduction stimuli. Hearing Research 181:131–143. 10.1016/S0378-5955(03)00183-7

[8] Allaire P, Raynor S, Billone M (1974) Cochlear partition stiffness—a composite beam model. The Journal of the Acoustical Society of America 55:1252–1258. 10.1121/1.1914693

[9] Steele CR (1974) Behavior of the basilar membrane with pure-tone excitation. The Journal of the Acoustical Society of America 55:148–162. 10.1121/1.1928144

[10] Rhode WS (1971) Observations of the Vibration of the Basilar Membrane in Squirrel Monkeys using the Mössbauer Technique. The Journal of the Acoustical Society of America 49:1218–1231. 10.1121/1.1912485

[11] Zhang X, Gan RZ (2011) A Comprehensive Model of Human Ear for Analysis of Implantable Hearing Devices. IEEE Transactions on Biomedical Engineering 58:3024–3027. 10.1109/TBME.2011.2159714

[12] Kim N, Steele CR, Puria S (2013) Superior-semicircular-canal dehiscence: Effects of location, shape, and size on sound conduction. Hear Res 301:72–84. 10.1016/j.heares.2013.03.008

[13] Kwacz M, Marek P, Borkowski P, Mrówka M (2013) A three-dimensional finite element model of round window membrane vibration before and after stapedotomy surgery. Biomech Model Mechanobiol 12:1243– 1261. 10.1007/s10237-013-0479-y

[14] Koike T, Sakamoto C, Sakashita T, et al (2012) Effects of a perilymphatic fistula on the passive vibration response of the basilar membrane. Hearing Research 283:117–125. 10.1016/j.heares.2011.10.006

[15] Borkowski P, Marek P, Niemczyk K, et al (2019) Bone conduction stimulation of the otic capsule: a finite element model of the temporal bone. Acta of Bioengineering and Biomechanics Vol. 21: 10.5277/ABB-01401-2019-02

[16] Secchia P, Cho N-H, McHugh CI, et al (2024) Towards motion measurements of the human organ of Corti using Optical Coherence Tomography (OCT). Mechanics of Hearing Workshop 2024 (MoH 2024), Ann Arbor, Michigan, USA. Zenodo.

[17] Viergever MA (1978) Basilar membrane motion in a spiral-shaped cochlea. The Journal of the Acoustical Society of America 64:1048–1053. 10.1121/1.382088

[18] Thorne M, Salt AN, DeMott JE, et al (1999) Cochlear Fluid Space Dimensions for Six Species Derived From Reconstructions of Three-Dimensional Magnetic Resonance Images. The Laryngoscope 109:1661–1668. 10.1097/00005537-199910000-00021

[19] Wysocki J (1999) Dimensions of the human vestibular and tympanic scalae. Hearing Research 135:39–46. 10.1016/S0378-5955(99)00088-X

[20] Merchant SN, Nadol Jr. JB (2010) Schuknecht’s Pathology of the Ear. PMPH-USA.

[21] Bhatt KA, Liberman MC, Nadol JB (2001) Morphometric Analysis of Age-Related Changes in the Human Basilar Membrane. Ann Otol Rhinol Laryngol 110:1147–1153. 10.1177/000348940111001212

[22] Plassmann W, Peetz W, Schmidt M (2008) The Cochlea in Gerbilline Rodents. Brain Behavior and Evolution 30:82–102. 10.1159/000118639

[23] Edge RM, Evans BN, Pearce M, et al (1998) Morphology of the unfixed cochlea. Hearing Research 124:1–16. 10.1016/S0378-5955(98)00090-2

[24] Cho NH, Wang H, Puria S (2022) Cochlear Fluid Spaces and Structures of the Gerbil High-Frequency Region Measured Using Optical Coherence Tomography (OCT). JARO 23:195–211. 10.1007/s10162-022-00836-4

[25] Dass R, Grewal BS, Thapar SP (1966) Human Stapes and its Variations: II. Footplate. The Journal of Laryngology & Otology 80:471–480. 10.1017/S0022215100065531

[26] Hato N, Stenfelt S, Goode RL (2003) Three-Dimensional Stapes Footplate Motion in Human Temporal Bones. Audiol Neurotol 8:140–152. 10.1159/000069475

[27] Singal A, Sahni D, Gupta T, et al (2020) Anatomic variability of oval window as pertaining to stapes surgery. Surg Radiol Anat 42:329–335. 10.1007/s00276-019-02347-z

[28] Sim JH, Röösli C, Chatzimichalis M, et al (2013) Characterization of Stapes Anatomy: Investigation of Human and Guinea Pig. JARO 14:159–173. 10.1007/s10162-012-0369-5

[29] Cohen D, Blinder G, Perez R, Raveh D (2005) Standardized computed tomographic imaging and dimensions of the round-window niche. Int Tinnitus J 11:158–162.

[30] Atturo F, Barbara M, Rask-Andersen H (2014) Is the Human Round Window Really Round? An Anatomic Study With Surgical Implications. Otology & Neurotology 35:1354. 10.1097/MAO.0000000000000332

[31] Singla A, Sahni D, Gupta A k., et al (2014) Surgical anatomy of round window and its implications for cochlear implantation. Clinical Anatomy 27:331–336. 10.1002/ca.22339

[32] Goycoolea MV, Lundman L (1997) Round window membrane. Structure function and permeability: A review. Microscopy Research and Technique 36:201–211. 10.1002/(SICI)1097-0029(19970201)36:3<201::AID-JEMT8>3.0.CO;2-R

[33] Guimarães CF, Gasperini L, Marques AP, Reis RL (2020) The stiffness of living tissues and its implications for tissue engineering. Nat Rev Mater 5:351–370. 10.1038/s41578-019-0169-1

[34] Miller CE (1985) Structural implications of basilar membrane compliance measurements. The Journal of the Acoustical Society of America 77:1465–1474. 10.1121/1.392041

[35] Kleppel MM, Santi PA, Cameron JD, et al (1989) Human tissue distribution of novel basement membrane collagen. Am J Pathol 134:813–825

[36] Dreiling FJ, Henson MM, Henson OW (2002) The presence and arrangement of type II collagen in the basilar membrane. Hearing Research 166:166–180. 10.1016/S0378-5955(02)00314-3

[37] Ishiyama A, Mowry SE, Lopez IA, Ishiyama G (2009) Immunohistochemical distribution of basement membrane proteins in the human inner ear from older subjects. Hearing Research 254:1–14. 10.1016/j.heares.2009.03.014

[38] Wenger MPE, Bozec L, Horton MA, Mesquida P (2007) Mechanical Properties of Collagen Fibrils. Biophysical Journal 93:1255–1263. 10.1529/biophysj.106.103192

[39] Kontomaris SV, Stylianou A, Malamou A (2022) Atomic Force Microscopy Nanoindentation Method on Collagen Fibrils. Materials 15:2477. 10.3390/ma15072477

[40] Kapuria S, Steele CR, Puria S (2017) Unraveling the mystery of hearing in gerbil and other rodents with an arch-beam model of the basilar membrane. Sci Rep 7:228. 10.1038/s41598-017-00114-x

[41] Motallebzadeh H, Soons JAM, Puria S (2018) Cochlear amplification and tuning depend on the cellular arrangement within the organ of Corti. Proceedings of the National Academy of Sciences 115:5762–5767. 10.1073/pnas.1720979115

[42] Maftoon N, Funnell WRJ, Daniel SJ, Decraemer WF (2015) Finite-Element Modelling of the Response of the Gerbil Middle Ear to Sound. JARO 16:547–567. 10.1007/s10162-015-0531-y

[43] Motallebzadeh H, Maftoon N, Pitaro J, et al (2017) Finite-Element Modelling of the Acoustic Input Admittance of the Newborn Ear Canal and Middle Ear. JARO 18:25–48. 10.1007/s10162-016-0587-3

[44] Gan RZ, Feng B, Sun Q (2004) Three-Dimensional Finite Element Modeling of Human Ear for Sound Transmission. Annals of Biomedical Engineering 32:847–859. 10.1023/B:ABME.0000030260.22737.53

[45] Homma K, Shimizu Y, Kim N, et al (2010) Effects of ear-canal pressurization on middle-ear bone- and air-conduction responses. Hearing Research 263:204–215. 10.1016/j.heares.2009.11.013

[46] Greenwood DD (1990) A cochlear frequency-position function for several species—29 years later. The Journal of the Acoustical Society of America 87:2592–2605. 10.1121/1.399052

[47] Johnstone BM, Patuzzi R, Yates GK (1986) Basilar membrane measurements and the travelling wave. Hearing Research 22:147–153. 10.1016/0378-5955(86)90090-0

[48] Robles L, Ruggero MA (2001) Mechanics of the Mammalian Cochlea. Physiological Reviews 81:1305–1352. 10.1152/physrev.2001.81.3.1305

[49] Allen JB (1980) Cochlear micromechanics—A physical model of transduction. The Journal of the Acoustical Society of America 68:1660–1670. 10.1121/1.385198

[50] Prodanovic S, Gracewski SM, Nam J-H (2019) Power Dissipation in the Cochlea Can Enhance Frequency Selectivity. Biophysical Journal 116:1362–1375. 10.1016/j.bpj.2019.02.022

[51] Prodanovic S, Gracewski S, Nam J-H (2015) Power Dissipation in the Subtectorial Space of the Mammalian Cochlea Is Modulated by Inner Hair Cell Stereocilia. Biophysical Journal 108:479–488. 10.1016/j.bpj.2014.12.027

[52] Perez-Flores MC, Verschooten E, Lee JH, et al (2022) Intrinsic mechanical sensitivity of mammalian auditory neurons as a contributor to sound-driven neural activity. eLife 11:e74948. 10.7554/eLife.74948

